# Time and space dimensions of gene dosage imbalance of aneuploidies revealed by single cell transcriptomes

**DOI:** 10.1101/424887

**Authors:** Georgios Stamoulis, Marco Garieri, Periklis Makrythanasis, Audrey Letourneau, Michel Guipponi, Nikolaos Panousis, Frédérique Sloan-Béna, Emilie Falconnet, Pascale Ribaux, Christelle Borel, Federico Santoni, Stylianos E Antonarakis

## Abstract

The mechanisms underlying cellular and organismal phenotypes due to copy number alterations (CNA) are not fully understood. Aneuploidy is a major source of gene dosage imbalance due to CNA and viable human trisomies are model disorders of altered gene expression. To understand the cellular impact of gene dosage imbalance, we studied gene and allele specific expression (ASE) of 9668 single-cell fibroblasts in trisomies T21, T18, T13 and T8. To limit the bias of interindividual noise, all comparisons between euploid and trisomic single-cells were performed on an isogenic setting for all trisomies studied. Initially we examined 928 single cells with deep RNA-Seq. For T21 we used fibroblasts from one pair of monozygotic twins discordant for T21 and from mosaic T21. For T18, T13 and T8 we analyzed single cells from mosaic individuals. Single-cell analyses revealed inconsistencies concerning the overexpression of some genes observed in differential trisomic vs euploid bulk RNAseq while this imbalance was not detectable in trisomic vs. euploid single cells. Moreover, ASE profiling of all single cells uncovered a substantial monoallelic pattern of expression in the trisomic fraction of the genome. By classifying genes according to the level of mono and bi-allelic transcription, we have observed that, for genes with monoallelic and low-to-average expression, the altered gene dosage is mainly due to the higher fraction of cells simultaneously expressing these genes in the trisomic samples. These results were confirmed in a further experiment of 8740 single fibroblasts from the monozygotic twins discordant for T21 samples. We conclude that gene dosage imbalance is of bidimensional nature: over time (simultaneous expression of all alleles resulting in increased accumulation of RNA of copy altered genes in each single cell) as previously stated, and over space (increased fraction of cells simultaneously expressing copy altered genes). These results strongly suggest that each class of genes contributes to the phenotypic variability of trisomies according to its temporal and spatial behavior and propose an improved model to understand the effects of copy number alterations.

## INTRODUCTION

The biochemical processes underlying complex cellular functions rely on a precise and timely dosage of their constitutive elements and, in particular, of protein stoichiometry^1^. Protein production is inherently connected with gene expression level, which, in turn, is regulated by several factors of genetic and epigenetic nature^2^. Perturbation of this equilibrium may induce severe cellular and organismal phenotypes. Genomic copy number alterations (CNA) such as duplications and deletions, result in gene expression imbalance^3^ and is associated with reproducible phenotypes as it is the case in aneuploidies. However the respective functional mechanisms are not well understood.

Aneuploidy is a well-known source of gene dosage imbalance through CNA. In particular trisomies are considered to be disorders of altered gene expression of the majority of genes on the supernumerary chromosomes (gene dosage sensitive genes)^4-8^. Trisomy 21 (T21 - Down syndrome) is the most common human aneuploidy compatible with postnatal survival, and has been extensively used as a model to study trisomies^9,10^. Other common trisomies include Trisomy 18 (T18 - Edwards’s syndrome) and Trisomy 13 (T13 - Patau syndrome)^11-14^. Phenotypes observed in trisomies have been attributed from bulk RNA-seq studies to the gene dosage imbalance, ~ 1.5 fold higher for the trisomic genes as compared to their euploid counterparts ^6,15-17^. However the causative links between altered gene expression and phenotypes in aneuploidies are not known. To understand the molecular basis of trisomy phenotypes, we explored gene expression profiles in single trisomic cells. Hitherto, single cell RNA-seq (scRNAseq) studies have revealed pervasive genome-wide skewed monoallelic gene expression in euploid cells ^18^, but also variability and gradation of gene expression for different genomic phenomena/processes, such as imprinting^19^ and X-inactivation^20^. Here we present the comparative analysis of scRNAseq from trisomic and matched isogenic euploid fibroblasts. We discovered that a significant component of gene dosage imbalance is actually explained by the significant fraction of trisomic cells that simultaneously activate gene expression as compared to euploid controls. These findings uncover a new dimension of gene dosage imbalance, constitutive of all regulatory mechanism of gene expression and copy number alterations and conceptually capable to explain the severity and variability of trisomy related phenotypes.

## RESULTS

### Accurate identification of trisomic cells in mosaic cell population

We used six different cell lines of skin fibroblasts from six individuals: two samples are from a pair of monozygotic twins discordant for T21^21^; four were from individuals mosaics for T21: CM05287, T13: GM00503, T18: AG13074, T8: GM02596 (Supplementary Figure 1 and Supplementary Table 1).

In order to classify trisomic and euploid cells in a mosaic trisomy cell population we developed an iterative clustering method based on k-means (k=2) using two metrics: the average cellular gene expression and ASE at heterozygous sites, both measured from the genes located on the supernumerary chromosome (details in Methods). Briefly, after quality control (doublets removal^20^ - Supplementary Figure 2) informative heterozygous sites were obtained by whole genome sequencing (WGS) and ASE was calculated considering the most covered heterozygous site per gene. Each site of the triplicated chromosome has the allele combination ABB or BAA with two identical alleles (double allele) and one unique allele. At each round of the iteration, the double allele is predicted for each heterozygous site and the status (euploid or trisomic) of a cell is (re)classified. Convergence is reached when the status of all cells is stable (Supplementary Fig 3A). We examined the accuracy of this method with a test-set of 316 euploid and trisomy single cells derived from a pair of monozygotic twins discordant for T21. We assigned the correct cellular status with an accuracy of ~95% (5-fold cross validation). As a further support of the reliability of the algorithm, the estimated fraction of trisomic cells in the different mosaic cell lines (mosaic T21, T8, T13, T18) was concordant with the degree of mosaicism derived by fluorescence in situ hybridization (FISH) (Supplementary Fig 3B). These results show that the single-cell ASE analysis in combination with the average cellular expression for triplicated genes can be used to computationally classify trisomic and euploid cells in samples from mosaic individuals.

### Random skewed monoallelic gene expression in trisomic single cell fibroblasts

Previous studies on allelic expression in euploid cell population have reported pervasive random skewed monoallelic gene expression at the single cell level ^18,22,23^, i.e cells expressed predominantly one allele (A or B) at a given time. We observed the same phenomenon in the trisomic cell population (Figure 1). More specifically for the euploid fraction of the genome, in twins’ fibroblasts discordant for T21, 60.1% of the heterozygous sites showed monoallelic expression (ASE ≤10.1; ASE≥0.9) in the euploid cells and 70.3% in T21 cells (Figure 1). Similar results were obtained for the euploid fraction of the genome for mosaic T21 cells and the other mosaic trisomies T8, T18 and T13 cells (Figure 2). In the monozygotic twins discordant for T21, the fraction of monoallelic ASE observations from chromosome 21 sites in euploid cells was 46.5% and 59.4% in T21 cells (Figure 1). In agreement with random selection of the transcribed allele, the fraction of trisomic informative sites exclusively expressing the unique allele (0≤ASE≤0.1) was ~1/3 of the total monoallelic observations. Accordingly, in all trisomy samples, the double alleles on the supernumerary chromosome were detected ~2 times more frequently than the unique alleles (Figure 2). Moreover, the mean of the distribution of biallelic observations (0.1<ASE<0.9) in trisomic single cells was not equal to 0.5 as in euploid single cells, but shifted towards 0.66 (Figure 1 and 2). These observations support a stochastic model of allelic selection by the transcriptional machinery where the probability of an allele to be expressed is linearly dependent on its respective copy number.

**Figure 1.**
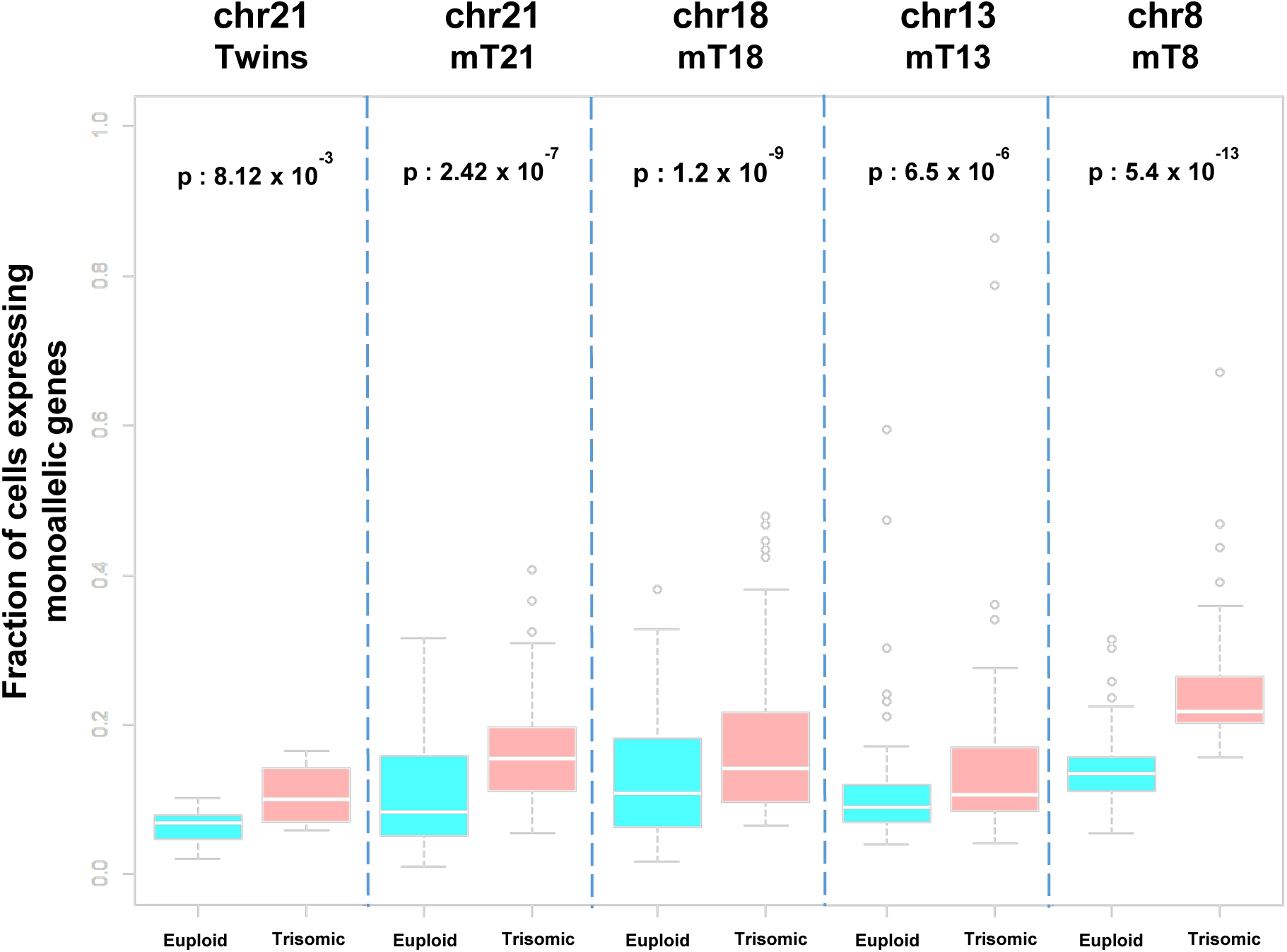
Histogram of SC ASE observations in Monozygotic Twins Discordant for DS. Upper panel. Histogram of genome wide ASE observations in SC excluding chr21 (Blue = Euploid Twin, Red = Trisomic Twin). High prevalence of monoallelic ASE observations were observed for both groups (60.14% Euploid – 70.3% Trisomic). Lower panel. Histogram of ASE observations in chr21 in single cells. Similarly to genome wide observations, monoallelic ASE in chr21 is prevalent (46.5% Euploid – 59.39% Trisomic. Notably, for trisomic SC, monoallelic ASEs on chr21 of the double allele (0.9-1) are twice as many of monoallelic ASEs of the single allele (0-0.1). Moreover biallelic observations in trisomic single cells are not centered at 0.5 as in euploid single cells, but at 0.66 (2/3).

**Figure 2.**
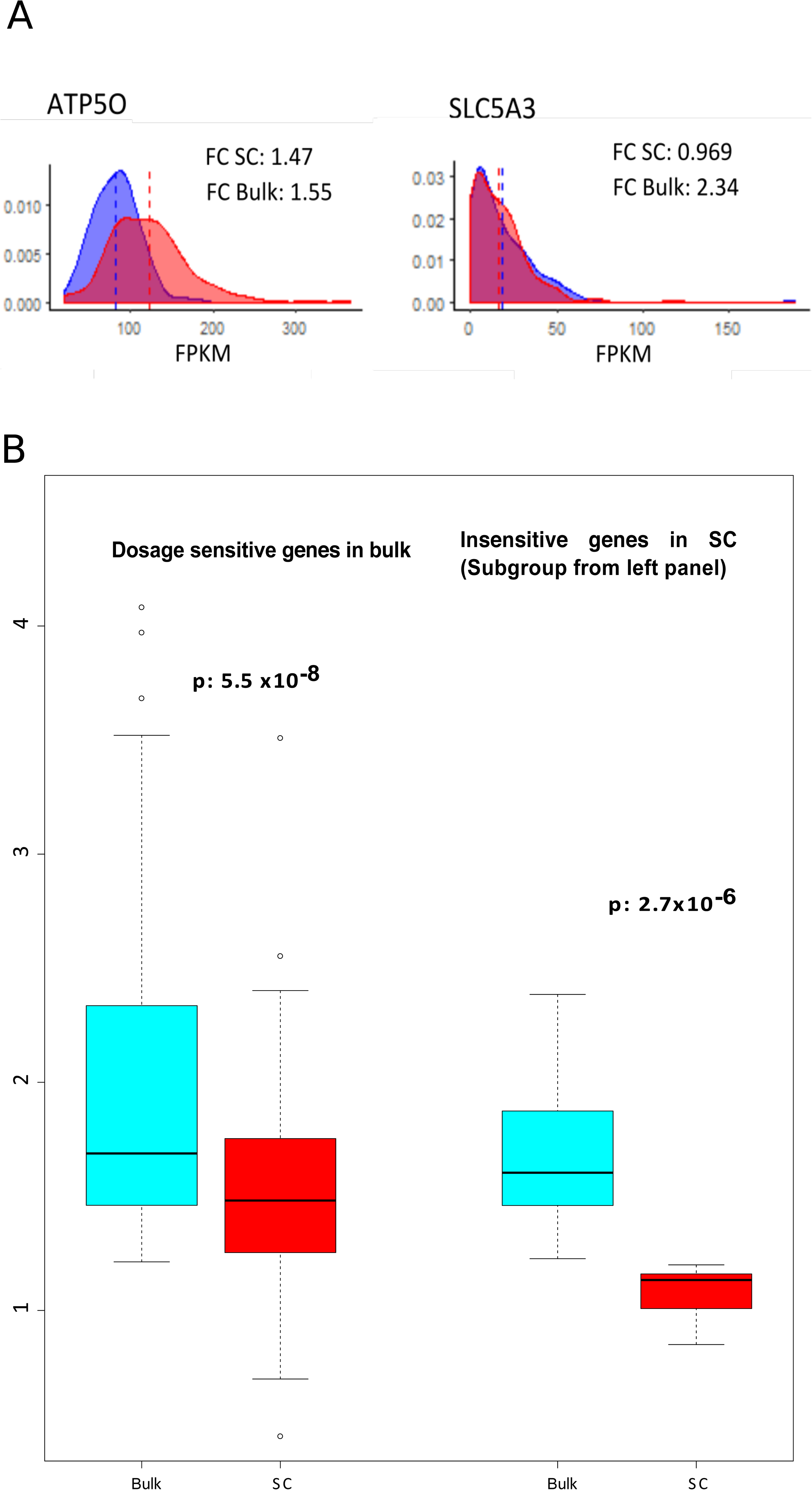
Histogram of SC ASE observations in individuals mosaic for different trisomies. Each row represents one mosaic individual. Left Panel: Histogram of genome wide ASE observations in SC excluding supernumerary chromosomes. High prevalence of monoallelic ASE observations was observed in all groups. Right panel: Histogram of ASE observations in SC in the supernumerary chromosomes. In all supernumerary chromosomes monoallelic ASE observations represent again the higher fraction of ASE observations, similarly to Genome Wide observations. In the Trisomic group, monoallelic observations on supernumerary chromosomes for double allele (0.9-1) are proportionally higher than monoallelic observations of the single allele (0-0.1). Moreover biallelic observations in Trisomic single cells are not centered as in euploid single cells, but shifted towards the double allele. (Blue = Euploid, Red = Trisomic)

### Monoallelic expression correlates with expression level

To further investigate, at the single cell level, the gene dosage effect in trisomic fibroblasts we classified the triplicated genes based on their monoallelic expression prevalence (MEP). MEP represents the fraction of cells per gene with ASE ≤10.1 or ASE≥0.9 at the heterozygous site with the highest number of cellular ASE observations (Supplementary Figure 4). We classified the triplicated genes in three groups: i) “Monoallelic” genes: MEP >80 % in euploid and trisomic cells; ii) “Intermediate” genes with MEP between 20% and 80%; and iii) “Biallelic” genes with MEP <20% (Figure 3A). Out of a total of 390 genes, 32% were classified as monoallelic, 66% as intermediate and only 2% as biallelic (Figure 3A, Supplementary Figure 5). Overall this classification was concordant for euploid and trisomic cells and consistent with previous single-cell RNA-seq study^24^.

**Figure 3.**
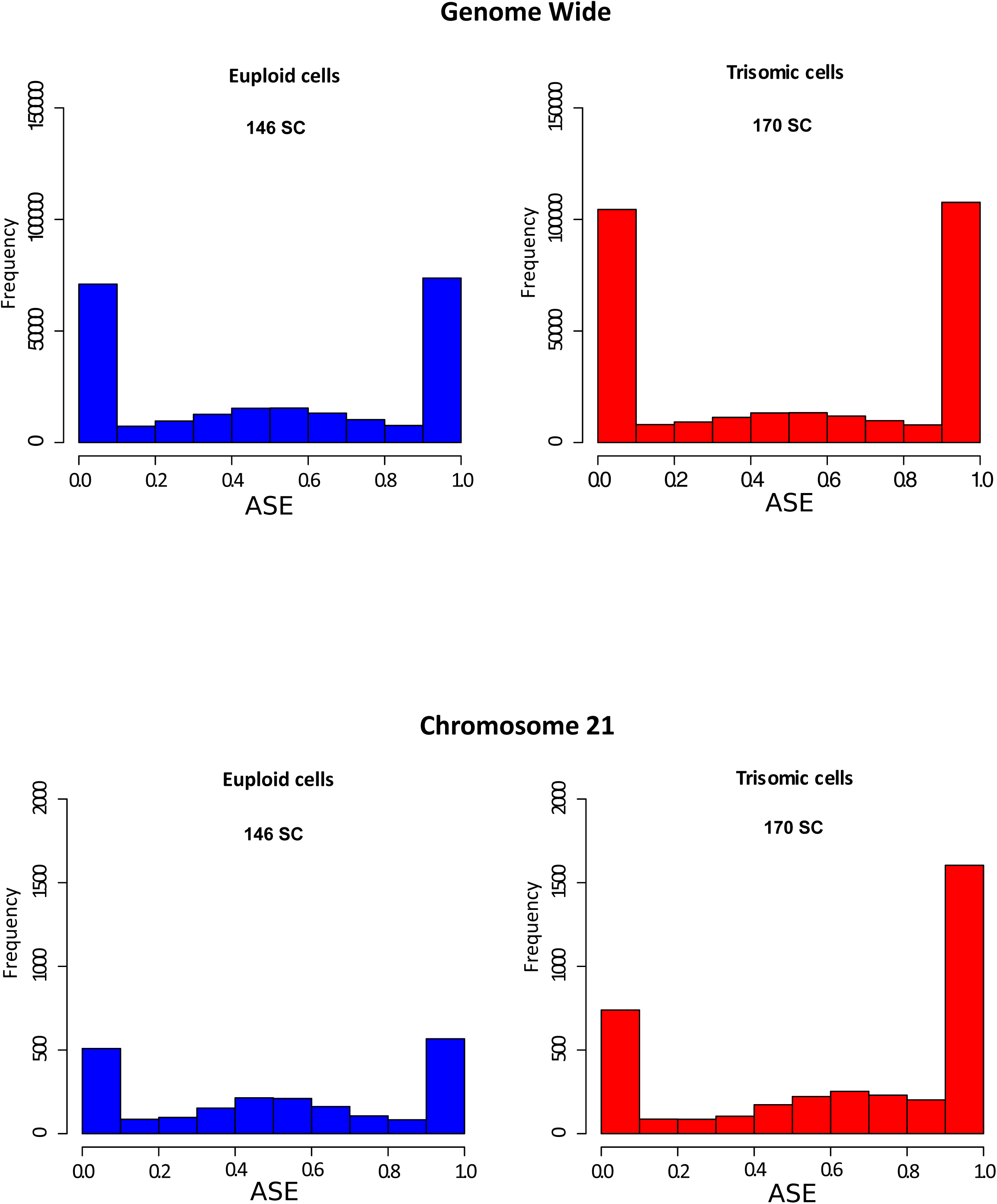
Prevalence of monoallelic expression in supernumerary chromosomes. A) Classification of genes in three groups (Monoallelic, Intermediate, and Biallelic). Genes for which > 80% of cells show monoallelic expression were classified as monoallelic (ASE ≤ 0.1 or ASE ≥ 0.9); genes with 20% to 80 % of cells with 0.1 ≤ ASE ≤ 0.9 were classified as intermediate; genes with <20% of cells with 0.1 ≤ ASE ≤ 0.9 were classified as biallelic. **B)** Monoallelic prevalence is negatively correlated with level of gene expression both genome wide (euploid fraction of the genome – upper panel) and within the triploid fraction of the genome (chr21 – lower panel).

We observed a significant negative correlation (ρ=-0.43, p=5e-3) between MEP and the level of gene expression of the corresponding triplicated gene (Figure 3B, Supplementary Figure 6). A similar negative correlation (ρ=-0.43, p<2.2e-16) was also observed for euploid genes in both the euploid and the trisomic cells (Figure 3B). According to the transcriptional bursting model of gene expression, highly expressed genes have short interburst periods and higher burst size (number of transcripts produced per burst) than low expressed genes^25^. Consequently it is frequent to observe in single cells random simultaneous (i.e. biallelic) transcription from the two alleles of highly expressed genes at a given time point. Conversely, low expressed genes have a low transcriptional bursting frequency and therefore the event of a biallelic simultaneous transcription is rare^22^.

### Effect of aneuploidy on gene expression is due to an increased number of expressor cells

In bulk studies, triplicated genes show overall the expected 1.5 gene expression fold change (trisomic vs euploid, FC). This observation has been attributed to the increased amount of transcripts produced by the triplicated genes. However our observation of extensive monoallelic expression of some of the triplicated genes in single cells challenges this interpretation of gene dosage imbalance in aneuploidies. As an example, the FC of SLC5A3 between normal and trisomic state is different between the bulk sample (FC=2.34) and across the single cells (meanFC=0.97) conversely, *ATP5O* has a FC of 1.55 in bulk and 1.47 in single cells. In general, we observed that dosage sensitive genes in the bulk have a significantly lower FC expression in single cells (Figure 4). FC for 94 chr21 dosage sensitive genes in the bulk sample is superior to 1.2 (T21/N) whereas many genes have a reduced or no gene dosage effect at the single cell level (Figure 4). We hypothesized a possible explanation of this phenomenon as follows.

**Figure 4.**
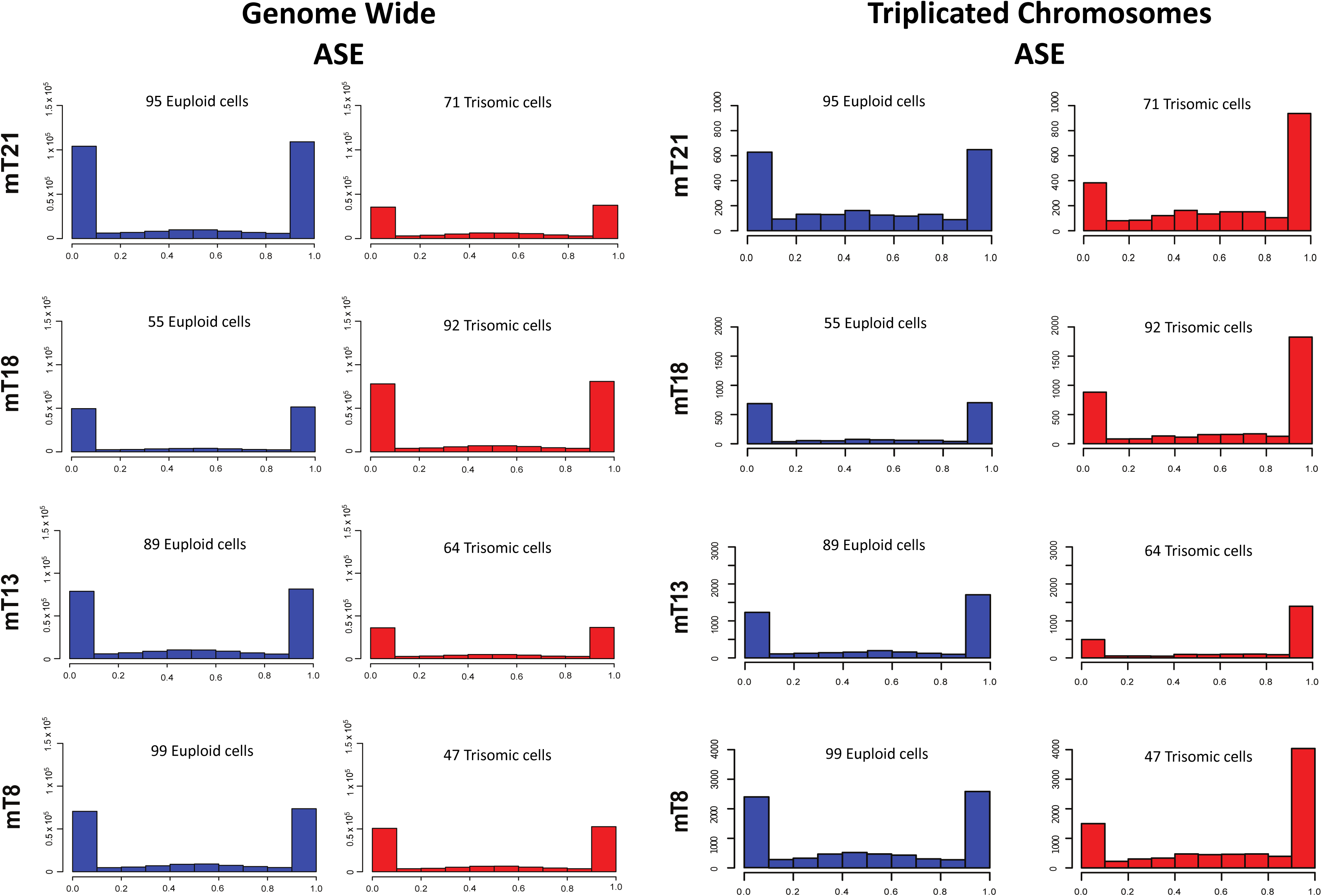
Fold change expression comparison in bulk and single cell study. A) Left: Distribution of expression levels of *ATP5O* in euploid (blue) and T21 (red) single cells. The gene presents with the typical trisomy gene dosage effect meanT21/mean =1.5 as observed in the bulk (FCbulk=1.5). Right: Distribution of expression levels of *SLC5A3* in euploid and T21 single cells. The two distributions are similar and the gene does not present the typical gene dosage effect as observed in the bulk (meanT21/meanD=1, FC_bulk_=2) B) Left: Comparison of expression fold change for dosage sensitive genes in the bulk (FC_bulk_>1.2, 94 genes) and SC of twins discordant for T21. Right: Comparison of expression fold change in bulk and SC for a subset of bulk-dosage sensitive genes presenting with a non-dosage sensitive effect in SCs (insensitive in SC) (0.8< SC FC<1.2, 17 genes).

Assuming the same total number of euploid and trisomic cells, the bulk expression of a gene g, E(g), can be decomposed in terms of the expression e of the gene g in each single cell of the bulk:

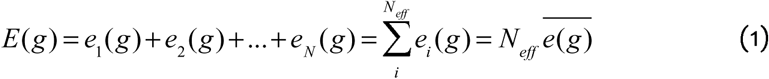

where N_eff_ is the number of cells expressing the gene g and 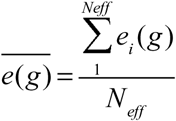 is the related mean expression. We can write the two equations for euploid (D) and trisomic (T) cells:

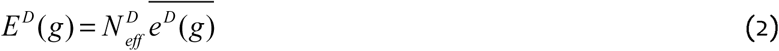

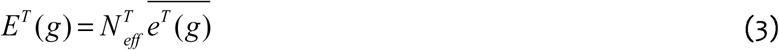

and the ratio of (3) vs (2) is

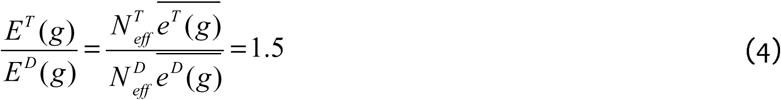

The final equivalence is given from the recurrently observed E^T^/E^D^ = 1.5 FC in trisomies. Equation (4) reveals the inverse proportionality between the mean FC in gene expression 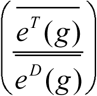 and the T/D ratio of the number of expressing cells 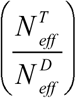.

More specifically (4) implies that genes in three copies with a similar expression level in single cells (i.e.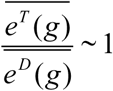) tend to be expressed in more trisomic cells than euploid cells 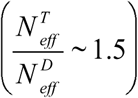. This theoretical result supports the general hypothesis that a component of bulk gene dosage imbalance of copy altered genes is generated by the increased number of cells expressing these genes at a given time point.

Along this hypothesis, we estimated the fraction of trisomic cells expressing each triplicated gene, defined as 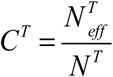 where N^T^ is the total number of cells in the trisomic sample and compared to the fraction of cells of the corresponding euploid sample 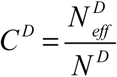. We consider a gene as expressed if: i) the total number of cells expressing the site within the gene is ≥20, ii) each cellular ASE observation has an RPSM score ≥20 (Reads Per Site Per Million), a metric we previously defined^18^. The genes were classified according to their prevalence of monoallelic expression (monoallelic, intermediate, biallelic) as previously defined. “Biallelic” or “intermediate” genes did not show statistically significant differences in the ratio of the fractions of expressing cells 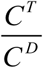. In contrast, in all trisomies, “monoallelic” genes present with a significantly higher fraction of cells expressing the triplicated genes than their respective euploid controls (Figure 5). Specifically, for the discordant monozygotic twins, the median fraction C^T^=10,8% vs C^D^=6.8% (8.1 × 10^−3^ paired Wilcoxon signed-rank test). For the mosaic T21 C^T^ =14.1% vs C^D^ =8.4% (2.4 × 10^−7^ paired Wilcoxon signed-rank test). All remaining mosaic samples (T18, T8, T13) showed a statistically significant higher fraction of expressing trisomic cells for “monoallelic” genes on the supernumerary chromosomes with respective p-values 1.2 × 10^−9^, 6.5 × 10^−6^, 5.5 × 10^−13^ (Figure 5).

**Figure 5.**
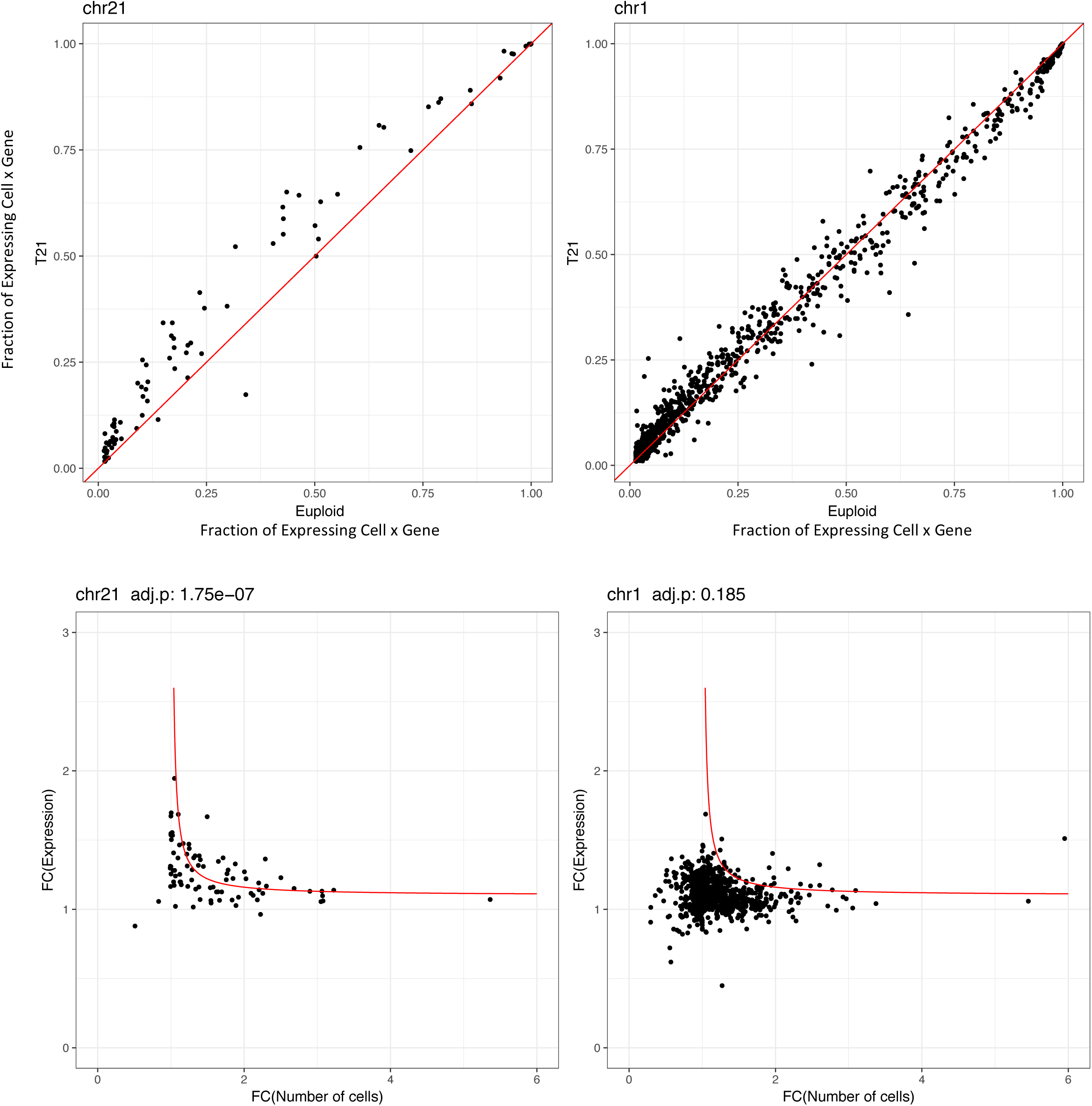
Higher fraction of expressed cells for supernumerary chromosome genes acts as an additional gene dosage mechanism in trisomies. In all trisomies an increased fraction of single trisomic cells expressing supernumerary chromosome monoallelic genes. Blue - fraction of euploid single cells. Red - fraction of trisomic single cells.

We validated these results with an additional and independent experiment using the chromium single cell controller (10X Genomics)^26^, a droplet-based system for scRNAseq. We processed 3801 euploid single cells and 4939 T21 single cells derived from the monozygotic twins fibroblasts discordant for T21. After random selection of an equal number of trisomic and euploid cells (3800) and normalization with respect to the total number of UMIs per cell, we confirmed that, only for chr21, FC expression of genes and respective FC of number of expressing cells fit the hyperbolic model of equation (4) (p=1×10^−7^, Spearman correlation). (Figure 6). As expected, for all genes in the autosomal chromosomes, the fraction of expressing cells was equivalent in both trisomic and euploid cells (Supplementary Figure 6). We concluded that this effect is exclusive of the supernumerary chromosome and thus likely implicated to gene dosage imbalance in T21. More specifically, we observed that gene dosage insensitive genes (0.8<FC<1.2), tend to exhibit a higher median fraction of trisomic vs euploid expressing cell ratio (1.3, p = 5 × 10^−10^) (Figure 7). This result points out the fraction of expressing cells as the main component of gene dosage imbalance for such genes. Notably, low expressed genes in chr21 (183 genes) showed a higher 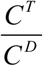 than intermediate (22 genes) and highly expressed genes (9 genes) (Figure 7). We conclude that for low expressed genes, the gene dosage imbalance is mainly driven by the higher fraction of T21 cells (p = 1×10^−6^ Figure 7). Conversely, for intermediate and highly expressed genes, the main component of gene dosage effect is the higher expression of triplicated genes in each single cell (p = 2×10^−5^ and p = 0.03 respectively, Figure 7).

**Figure 6.**
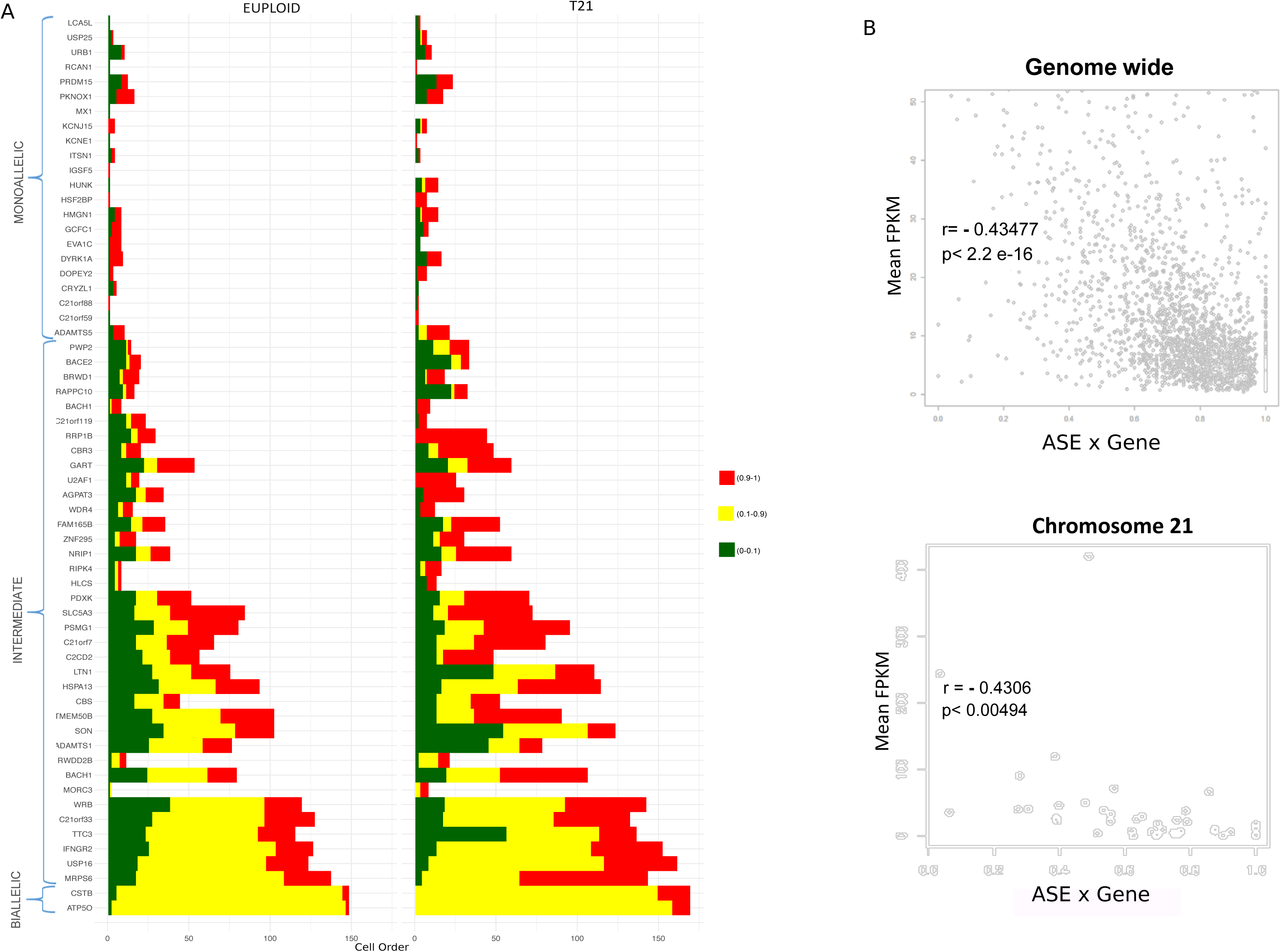
Validation of higher fraction of expressing cells of trisomic genes in 8740 single fibroblasts. Upper row, left: (y-axis) distribution of fraction of trisomic cells expressing chr21 genes; (x-axis) distribution of fraction of euploid single cells expressing chr21 genes; right: data for chr1 as control. Lower row, left: T/D ratio of number of expressing cells and T/D ratio of single cell expression of genes in chr21 are inversely correlated (Spearman correlation); right: data for chr1 as control. Cells with >5 reads and genes expressed in >50 cells have been considered. Red line is to guide the eye (see text for details).

**Figure 7.**
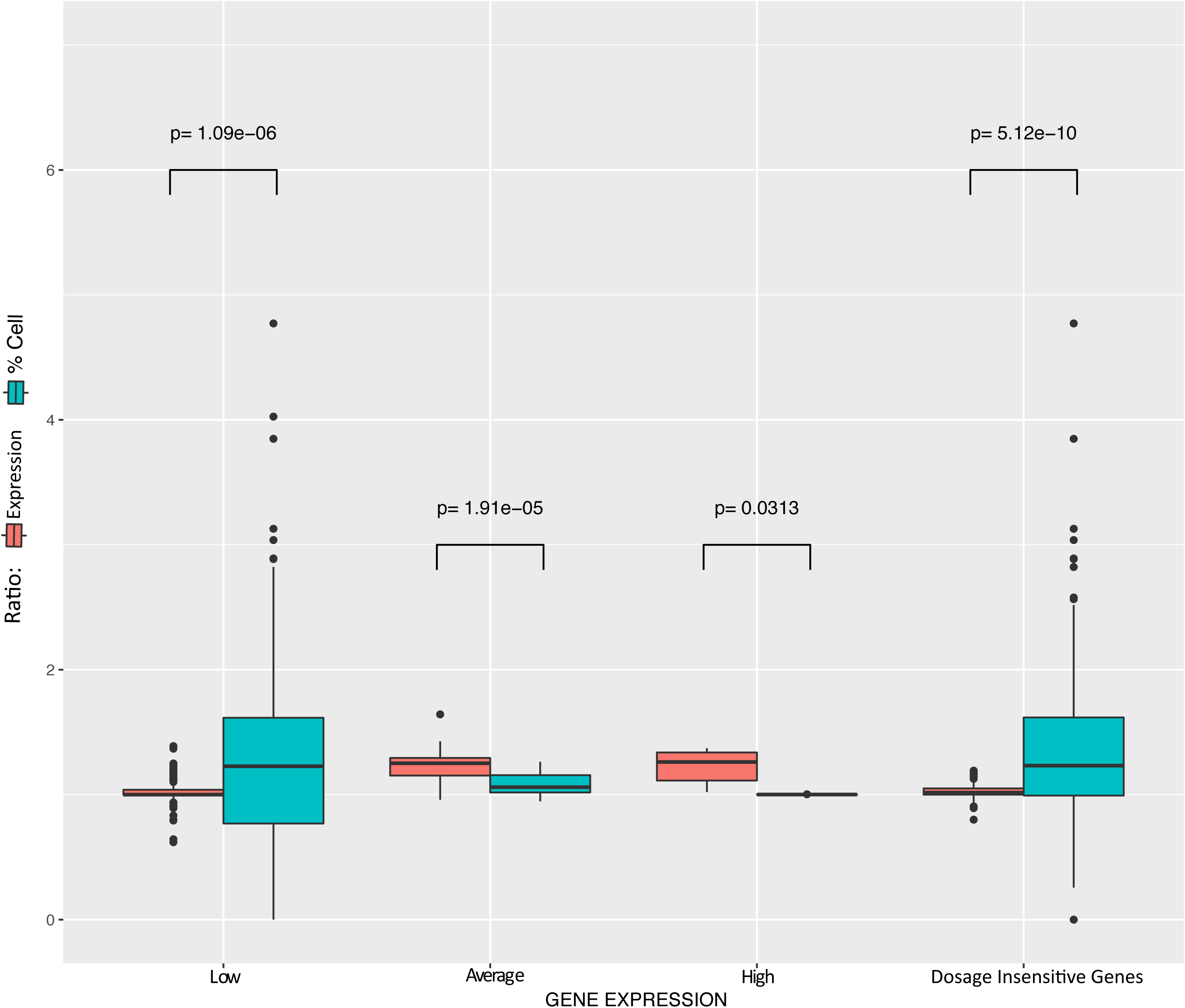
Components of gene dosage imbalance in trisomy 21 using 8740 single fibroblasts. Left: Gene dosage imbalance time and spatial components in low (<3 FPKM), medium (3 FPKM< and <15 FPKM) and high (>15 FPKM) expressed genes. For low expressed genes dosage imbalance is mainly driven by the increased fraction of trisomic cells expressing these genes compared to euploid. For medium and highly expressed genes the dosage imbalance is mainly driven by Trisomic/Euploid FC expression per cell while no significant difference in the fraction of cells can be detected. Right: statistically significant differences of Trisomic/Euploid ratio of fraction of expressing cells in non dosage sensitive genes in single cells (0.8<FC<1.2).

## DISCUSSION

Gene dosage imbalance caused by CNA is generally interpreted as the increased production of transcripts per cell. The emblematic example is represented by trisomies where several studies on mouse and human samples reported the classical 1.5 average gene expression fold change^27-35^. The results of our study strongly suggest a more complex scenario. As a first striking observation, we detected reduced or no dosage effect at all for some genes on a single cell level, compared to the expected fold change from our previous bulk study in T21. Moreover, ASE exploration of the supernumerary chromosome genes in our isogenic models of trisomies showed clear random monoallelic patterns of expression as already observed in euploid cells^18,23,24^. We confirmed that these patterns follow a random allelic selection model by observing that the number of observations expressing the duplicated allele was indeed twice the number of those expressing the single allele. An additional observation of this study indicated that the monoallelic prevalence of expression (the fraction of cells in which the gene appears as expressed by one allele only) is negatively correlated with the level of expression of the respective genes.

Finally, we observed that the increased fraction of trisomic cells vs. euploid presenting with active expression of supernumerary chromosome genes is contributing to the average dosage imbalance of all the trisomies analyzed in this study. This effect is more evident for low expressed/monoallelic genes. Taken together these results suggest that the presence of the extra chromosome and its availability to the transcription machinery increase the probability of transcription of the duplicated genes in time (more alleles simultaneously expressed in one cell) and in space (more cells expressing the same genes at a given time point) with respect to euploid controls. For average and highly expressed genes (i.e. maintenance of cell function), transcriptional bursting events are frequent in both euploid and trisomic single cells. Accordingly, we observed no significant difference in the fraction of expressing cells. We concluded that the dosage effect for these genes is mainly of temporal nature. Conversely the dosage effect for low expressed genes, characterized by random monoallelic expression, is mainly of spatial nature thus caused by the higher fraction of expressing cells.

This observation may have a significant impact on the understanding the molecular pathophysiology of aneuploidies. “Spatial” gene dosage imbalance in single cells could be crucial on a tissue level in the following (the list is not exhaustive): 1) different fractions of cells producing increased level of subunits of multimeric proteins may result in abnormal stoichiometry^36^; 2) abnormal number of cells expressing fundamental transcription factors^37^; 3) abnormal number of cells with cell surface receptors and ligands that may results in a disturbed developmental fate^38,39^; 4) abnormal number of transporter molecules in the tissue resulting in metabolic disturbances^40^; 5) excess of cell adhesion molecules that may increase cellular adhesiveness and differential fate of a tissue^41^; 6) alteration in the production, concentration and diffusion of morphogens in the tissue and consequent abnormal cellular proliferation and development of aberrant cellular and tissue structures^42^. Furthermore, the unbalanced expression of long non-coding RNAs and microRNAs in a fraction of cells may also contribute to the disturbance of the regulatory repertoire of other cells, particularly during embryogenesis^43^. Indeed many of these phenotypes may manifest during the early embryonic development stages where a precise and delicate balance among gene pathways dedicated to coordinate cell-to-cell interactions must be maintained^44^. Additionally this effect can be mediated by the duplication of regulatory regions that modulate gene expression through specific regulatory variants. eQTLs in trisomic regions have 4 possible states (AAA, AAB, ABB, BBB) instead of the canonical (AA, AB, BB) in the euploid genome. This additional degree of freedom, and the two dimensions of gene dosage imbalance, might contribute to the considerable phenotypic variability among affected individuals. More generally, we propose that spatial gene dosage may contribute to phenotypes related to Copy Number Alteration, including Copy Number Variants (CNVs) and somatic partial aneuploidies typical of cancer cells^45^. Time-series single cell RNA-seq studies in aneuploid embryos are needed to reveal how time and spatial dimensions of gene dosage imbalance interplay to determine individual phenotypic features.

## MATERIAL AND METHODS

### Ethical statement

The study was approved by the ethics committee of the University Hospitals of Geneva, and written informed consent was obtained from both parents of each individual.

### Samples

We used six different cell lines of skin fibroblasts from six individuals: two samples are from a pair of monozygotic twins discordant for T21^21^; four were from individuals mosaics for T21: CM05287, T13: GM00503, T18: AG13074, T8: GM02596 (https://www.coriell.org/). DNA samples from peripheral blood were obtained from the parents of the monozygotic twins. Cell lines from mosaic individuals T8, mosaic T13, mosaic T18, were purchased from Corriel, and sample from mosaic T21 was kindly provided by Prof. Dean Nizetic. We captured in total 928 single-cell fibroblasts (484 Euploid and 444 Trisomic) using the Fluidigm C1 technology. In addition we employed an alternative single cell RNA-seq protocol based on 10X Genomics technology (Chromium Single cell 3’ Solution protocol^26^) to capture 8740 single cells (3801 euploid and 4939 trisomic single cells) **(Supplementary Table 1).**

### Analysis of genome-matched samples

The comparison of transcriptional profiles of unrelated individuals is complicated by the substantial genetic variability ^34^. Notably, in this study, we sought to eliminate the inter-individual bias by comparing euploid and trisomic single cell fibroblasts from individuals with mosaicism for the relevant trisomies (T8, T13, T18, T21) and by using single cell fibroblasts from monozygotic twins discordant for DS (T21) (Supplementary Figure 1 and Supplementary Table 1).

### Cell Culture

Cells were cultured in DMEM GIutaMAX^™^ (Life Technologies) supplemented with 10% fetal bovine serum (Life Technologies) and 1% penicillin/streptomycin/fungizone mix (Amimed, BioConcept) at 37°C in a 5% CO2 atmosphere. The day before the single-cell capture experiment; cells were trypsinized (Trypsin 0.05%-EDTA, Life Technologies) and replated at a density of 0.3 × 10^6^ cells/100-mm dish.

### Fluorescence in situ Hybridization

Fluorescent in situ hybridization (FISH) analysis was performed on cultured interphase nuclei with 2 set of probes including two locus specific probes on chromosome 13 (Vysis, RB1;13q14 locus) and chromosome 21 (Vysis, D21S342/D21S341/D21S259 contig probes) for set 1 and two alpha satellite centromere probes for chromosome 8 (Vysis, D8Z1) and chromosome 18 (D18Z1) for set 2. The experiments were carried out according to manufacturer’s instructions (Aneuvysion, VYSIS, Inc). For each sample 100 interphase nuclei were examined to evaluate mosaics rate.

### Whole Genome Sequencing

Genomic DNA extraction was performed using a QIAamp DNA Blood Mini Kit (Qiagen). DNA was fragmented by Covaris to sizes of ~300 bp. Single cell libraries preparation was performed with TruSeq DNA kit (Illumina) with input of 1 μg of gDNA and sequenced on an Illumina HiSeq 2000 system in paired end reads of 2×100-bp as previously described^18^. All experiments procedures were followed according to manufacturer’s instructions and protocols. For each individual, whole genome DNA sequencing was analyzed using an in-house pipeline, using BWA^46^ for mapping the reads over the hg19 reference genome, SAMtools^47^ for detection of heterozygous sites, and ANNOVAR for the annotation of variants^48^. For the monozygotic twins discordant for DS, the assignment and reconstruction of the haplotypes were done using genotyping data from the parents. The double allele was derived from the mother. Using ShapeIT^49^ we phased the haplotypes for the single and double allele of chr21.

### Single-cell capture (C1 Fluidigm)

Single-cell capture was performed by C1 single-cell auto prep system (Fluidigm) following the manufacturer’s instructions^18^. The microfluidics circuit used was the C1^™^ Single-Cell mRNA-seq IFC, 17–25 μm. All 96 chambers were inspected under an inverted phase contrast microscope; only chambers containing a non-damaged single cell were considered for downstream analysis. For the cell lysis and cDNA synthesis, we used the SMARTer Ultra Low RNA kit for Illumina Sequencing (version 2, Clontech) and a C1 Auto Prep System instrument (Fluidigm) with the original mRNA Seq Prep script provided by the manufacturer (1772×/1773×, Fluidigm). We assessed cDNA quality on 2100 Bioanalyzer (Agilent) with the high sensitivity DNA chips (Agilent) and quantified the cDNA using Qubit dsDNA BR assay kit (Invitrogen). Sequencing libraries were prepared with 0.3 ng of pre-amplified cDNA using Nextera XT DNA kit (Illumina) according to manufacturer’s instructions. Libraries were sequenced on an Illumina HiSeq2000 machine as 100 bp reads single-end.

### GemCode single-cell libraries preparation and sequencing

We captured in total 3801 euploid and 4939 trisomic single cell fibroblasts from the monozygotic twin pair using the Chromium System powered by GemCode Technology (10× Genomics)^26^. Single-cell RNA-seq libraries were generated using the Chromium Single Cell 3’ Reagent Kit version 2 (10× Genomics) according to the manufacturer’s instructions. Briefly, the concentration of trypsin dissociated fibroblasts was set to 1500 cells/μl of culture medium (Dulbecco’s Modified Eagle Medium (DMEM), 10% FBS) and 5000 individualized cells were flown per channel following the recommendation of the manufacturer. All libraries were quantified by Qubit (Invitrogen) and by quantitative real-time PCR using the PCR-based KAPA Library Quantification Kits for Illumina platforms (Kapabiosystems). Size profiles of the preamplified cDNA and sequencing libraries were assessed using a 2100 BioAnalyzer (Agilent) with a High Sensitivity DNA chip kit (Agilent). Barcoded libraries were sequenced with an HiSeq 4000 (Illumina) as paired-end 100 bp reads as recommended by 10× Genomics.

### C1 Single-cell RNA-sequencing

For single cells capture with the Fluidigm C1 microfluidics system, SMARTer Ultra Low RNA kit for Illumina sequencing (version 2, Clontech) was used for cell lysis and cDNA synthesis. 0.3 ng of pre-amplified cDNA, was used for the library preparation with the Nextera XT DNA kit (Illumina) as described^18^. Libraries were sequenced on an Illumina HiSeq2000 sequencer as 100 bp single-ended reads. RNA sequences were mapped with GEM^50^. Uniquely mapping reads were extracted by filtering for mapping quality (MQ>=150). For FPKM expression quantification an in-house algorithm was used with GENCODE V19 as reference. Cells with less than 10 million reads and/or cells with <10% of expressed genes (total number of 56680 genes)

### Allele-specific expression

Cellular Allelic Specific Expression (ASE) of each heterozygous site was calculated in the euploid and triploid fraction of the genome of each single cell per individual using two different formulas.

Euploid genome:

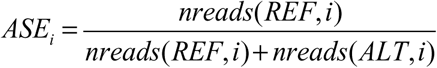

Triploid fraction:

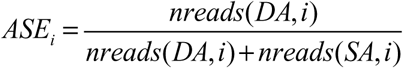

where nreads is an operator giving the number of reads covering the site i, mapped according to the REFerence or the ALTernative allele (euploid) or to the Double or Single Allele (triploid).

In both cases, ASE values range from 0 to 1 (Supplementary Figure 2). We consider 0≤ASE≤0.1 as the signature of monoallelic expression of the Alternative allele (euploid) or Single allele (triploid). Conversely 0.9≤ASE≤1 indicates monoallelic expression of the Reference allele in the case of euploid cells or of the Double allele in the case of (trisomic cells). ASE from 0.1 to 0.9 is an indicator of biallelic expression.

### Identification of Euploid and Trisomic single cells of mosaic populations

We developed a computational procedure to distinguish euploid from trisomic single cells in mosaic populations. Using an iterated k-means (k=2) approach we combined ASE profiling and expression data from the supernumerary chromosomes to classify each single cell as euploid or trisomic. The main idea is to estimate for each heterozygous site the allele in two copies (allele with REF or ALT genotype) using ASE imbalance of single cells for each trisomy studied (ASE≥0.65 implies REF allele in two copies, ASE <0.65 implies ALT allele in two copies). Once this estimation is done, the phased ASE can be used as a second cluster dimension to discriminate trisomic from euploid cells (the first dimension is the average gene expression, supposedly higher in trisomic cells). At the first step the double allele is randomly defined for each site and cells are clustered as trisomic or euploid with respect to the average gene expression. With this first classification, the algorithm refines the estimation of the double allele in the new set of trisomic cells only, according to average ASE (across all trisomic cells) per site. K-means cell clustering and allele estimation are repeated until convergence is reached (i.e. trisomic and euploid cell clusters are stable, no reassignment)(Supplementary Figure 3).

### Fluidigm C1 multiple cells (doublets) detection

In our Fluidigm C1 based protocol, we set two checkpoints where double cells (doublets) are identified and eliminated. First, during the capturing procedure, doublets are identified by visual inspection under the microscope, and eliminated from further analysis. Second, after RNA sequencing and ASE analysis, potential double cells of female individuals are eliminated based on the study of X chromosome haplotype expression. For each cell, the expressed haplotype is estimated by calculating the allelic ratio of each heterozygous site in the X chromosome as identified by whole genome sequencing. Sites in the pseudoautosomal regions (PAR1 chrX:60001-2699520, PAR2 chrX:154931044-155260560) and known escapee genes are *a priori* excluded. The estimated haplotype of each cell was compared to all the others through correlation based hierarchical clustering. Cells expressing concordant and discordant haplotypes results in a correlation near 1 and −1 respectively. Doublets simultaneously expressing both discordant haplotypes cluster around the absolute correlation of 0.5 and are excluded from further analysis (Supplementary Figure 2).

### Allele drop out control

To reduce the potential bias induced by allele drop-out, we have previously defined the RPSM metric (Reads per site per million mapped reads)^18^. Through split cell RNA experiments based on ERCC RNA spike-in mix (Ambion)^18^, we identified the threshold RPSM=20 to drastically reduce false positive monoallelic ASE calls (Supplementary figure 7). Additionally we only consider heterozygous SNV sites covered by at least 16 reads to further minimize possible allele drop out effects^51^.

